# Methylation at nucleotide C^62^ in spliceosomal RNA U6 alters mRNA splicing which is important for embryonic development

**DOI:** 10.1101/661876

**Authors:** Allison Ogren, Nataliya Kibiryeva, Jennifer Marshall, James E. O’Brien, Douglas C. Bittel

**Affiliations:** College of Biosciences, Kansas City University of Medicine and Biosciences (KCU), Kansas City, MO, USA; Ward Family Heart Center, Children’s Mercy Hospital, Kansas City, MO, USA

**Author notes:** (DCB).

## Abstract

Understanding the regulation of development can help elucidate the pathogenesis behind many developmental defects found in humans and other vertebrates. Evidence has shown that alternative splicing of messenger RNA (mRNA) plays a role in developmental regulation, but our knowledge of the underlying mechanisms that regulate alternative splicing are inadequate. Notably, a subset of small noncoding RNAs known as scaRNAs (small cajal body associated RNAs) contribute to spliceosome maturation and function through covalently modifying spliceosomal RNAs by either methylating or pseudouridylating specific nucleotides, but the developmental significance of these modifications is not well understood. Our focus is on one such scaRNA, known as SNORD94 or U94, that methylates one specific cytosine (C^62^) on spliceosomal RNA U6, thus potentially altering spliceosome function during embryogenesis. We previously showed that mRNA splicing is significantly different in myocardium from infants with congenital heart defects (CHD) compared to controls. Furthermore, we showed that modifying expression of scaRNAs alters mRNA splicing in human cells, and zebrafish embryos. Here we present evidence that SNORD94 levels directly influence levels of methylation at C^62^ in U6, which we have previously shown is associated with altered splicing and congenital heart defects. The potential importance of scaRNAs as a developmentally important regulatory mechanism controlling alternative splicing of mRNA is unappreciated and needs more research.

**Author summary:** Splicing of mRNA transcripts by removal of introns and some non-critical exons is a crucial part of mRNA processing, gene expression, and cell function, and regulation of this process is still under investigation. Alternative splicing of mRNA transcripts of genes is tissue and time specific throughout life, although this process occurs everywhere in the body according to local tissue needs and signals. The spliceosome, the large ribonucleoprotein complex that carries out splicing, is biochemically modified by small noncoding RNAs, which is important for its structure and function. Here we show that the amount of 2’-O-ribose methylation at nucleotide C^62^ in spliceosomal RNA U6 is dependent on the level of the scaRNA SNORD94. We hypothesize that alternative splicing is dependent, at least in part, on biochemical modification to the spliceosomal RNAs. Furthermore, when scaRNA directed modifications are dysregulated, the result causes inappropriate alternative splicing that may contribute to developmental defects such as congenital heart defects. To our knowledge, this is the first demonstration that 2’-O-ribose methylation is indeed dependent on scaRNA levels in human cells and tissues.

## Introduction

Congenital heart defects are structural problems in the heart that are present at birth that affect normal function and blood flow through the heart [1]. Nearly 1% of children are born with a congenital heart defect (CHD) every year, making it the most common type of birth defect [1]. Despite how common these defects are, a resounding majority (>70%) of CHDs have unknown etiology to date [2]. The most common complex CHD is tetralogy of Fallot (TOF), which is a combination of four defects and has an incidence of 5 to 7 in every 10,000 live births (5% to 7% of all congenital heart lesions) [1, 3]. TOF includes pulmonary stenosis, a large ventricular septal defect, an overriding aorta, and right ventricular hypertrophy. The result of this condition is oxygen-poor blood from the right ventricle being pumped into the aorta rather than the pulmonary artery, and an overworking of the right ventricle that causes the hypertrophy [1]. TOF is treated with surgical intervention, the first of which occurs within the first year of life with potential further or repeated surgeries later in life.

Chromosomal defects and single gene disorders are known to cause some CHDs, often in the context of multisystem diseases [2]. Genetic mechanisms underlying nonchromosomal or non-Mendelian “sporadic” CHD are poorly understood although these sporadic cases account for the majority of CHDs [3]. Occurrence of CHD in children of mothers with TOF is approximately 3.1%, which is significantly higher than the overall occurrence, supporting a genetic contribution; however, the exact basis is not yet understood [3]. Children with sporadic CHD are most often born to unaffected parents, suggesting incomplete penetrance which can be a consequence of differences in genetic buffering capacity between individuals [3-5]. Also, de novo events, including sequence alteration or copy number changes, can have an impact on gene function or alter dosage and contribute to mutational load [3]. Additionally, recessive mutations, when homozygous, can further destabilize regulatory networks [3]. This makes identification of human disease genes involved in sporadic CHD by classical positional genetics very difficult, and so other approaches are needed.

During embryonic development, spatiotemporal signaling between the first and second heart field gives rise to the left and right ventricles and conotruncal outflow tract, which is critical for correct development of the vertebrate heart [6]. The genetic factors implicated in sporadic CHD have been reviewed previously, but these known genetic factors still account for a small percentage of TOF cases [7]. We examined alternative splicing of the transcriptome in heart tissue from infants with TOF. We focused on important regulatory genes involved in heart development in particular (*GATA4, MBNL1, MBNL2, DICER, DAAM1*, and *NOTCH2*), and showed that the alternative splice isoforms of these genes were abnormal in TOF patients compared to control, and they were in fact more similar to fetal tissue splice isoforms [8]. We hypothesized that regulatory pathways may be disrupted by alternative splicing, contributing to developmental disorganization.

Many human diseases are the result of aberrant pre-mRNA splicing and therefore understanding splicing on a molecular level is of medical relevance [9]. Several studies have shown that mRNA splicing is highly dynamic and intricately regulated during vertebrate heart development [10-13]. Particularly, the transition from fetal to postnatal patterns of a conserved set of alternatively spliced isoforms has been shown to be critical for proper mouse heart development [14]. It is apparent that alternative mRNA splicing plays a crucial role in heart development, but the role in human heart pathology is not yet clear. Recently, evidence has begun to mount that spliceosome maturation processes contributing to splicing fidelity may play a critical role in mRNA isoform transitions during vertebrate heart development [6].

The spliceosome is a multimegalodalton ribonucleoprotein complex comprised of several subunits (U1, U2, U4, U5, and U6) that carries out specific mRNA processing including splicing of introns and exons according to the specific needs of the cell [9]. During spliceosome maturation, the spliceosomal RNAs (snRNAs) are biochemically modified by a set of small noncoding RNAs, the scaRNAs (small cajal body-associated RNAs) [8]. The scaRNAs are a subset of the small nucleolar RNA (snoRNA), which is a large family of noncoding RNA that is known to primarily guide biochemical modifications on specific nucleotides of rRNAs and snRNAs. It has been shown that without the specific modifications controlled by the scaRNAs, the spliceosome fails to function properly [15]. Two distinct families of snoRNA exist (C/D box and H/ACA box), based on conserved nucleotide motifs [16]. The C/D box family facilitates 2’-O-ribose methylation on spliceosomal snRNAs, and the H/ACA box facilitates pseudouridylation, with a few exceptions [16].

The 2’-O-methylated sites tend to reside in functionally important regions in ribosomal RNA, potentially influencing ribosomal structure and function [17]. Within spliceosomal RNA, the methylated nucleotides present are also 2’-O-methylated, suggesting a similar structural and functional role within the spliceosome to these methyl groups in the ribosome [15, 16]. Within spliceosomal RNA U6, there are 3 examples of 2’-O-methylcytidine, one of which is the target of SNORD94 (previously called U94), and the other two are targets of SNORD67 and SNORD10 [18]. For the purpose of this study, we will focus on SNORD94, which is a C/D box scaRNA and targets a cytosine at position 62 (C^62^)on spliceosomal RNA (snRNA) U6 for 2’-O-methylation [19]. The 2’-O-methylation at C^62^ on snRNA U6 lies within the stem-loop structure of the catalytic site of the activated U6 RNP, potentially contributing to the catalytic site structure and function and influencing the spliceosome’s ability to splice correctly.

Previously, it has been shown that snoRNA expression is different in the right ventricular tissue of children with TOF, with 135 snoRNAs being significantly differently expressed from control tissue, most of which having decreased expression in TOF samples [3]. In fact, a similarity in expression levels of most snoRNAs between children with TOF and fetal tissue has been shown, with 115 out of 126 snoRNAs with decreased expression and 6 out of 9 with increased expression relative to control tissue [3]. Of those snoRNAs with decreased expression in patients with TOF, 6 targeted 5 nucleotides (of 8 total) on U6 according to the snoRNA database [3, 18]. SNORD94 was among these snoRNAs with reduced expression in children with TOF as well as fetal tissue relative to right ventricular tissue from normally developing infants. We hypothesize that SNORD94 expression level along with the other scaRNAs is important for spliceosomal function and may therefore may play a role in regulating heart development.

Of note, there are other known sites of 2’-O-ribose methylation within spliceosomal RNA U6 (Figure 1). The scaRNAs that modify C^60^, HBII-166 (SNORD67), has been shown to have reduced expression in RV tissue from infants with TOF, but the scaRNA that modifies C^77^, mgU6-77 (SNORD10), is not known to have altered expression in RV tissue from infants with TOF[3, 8].

**Figure 1.**
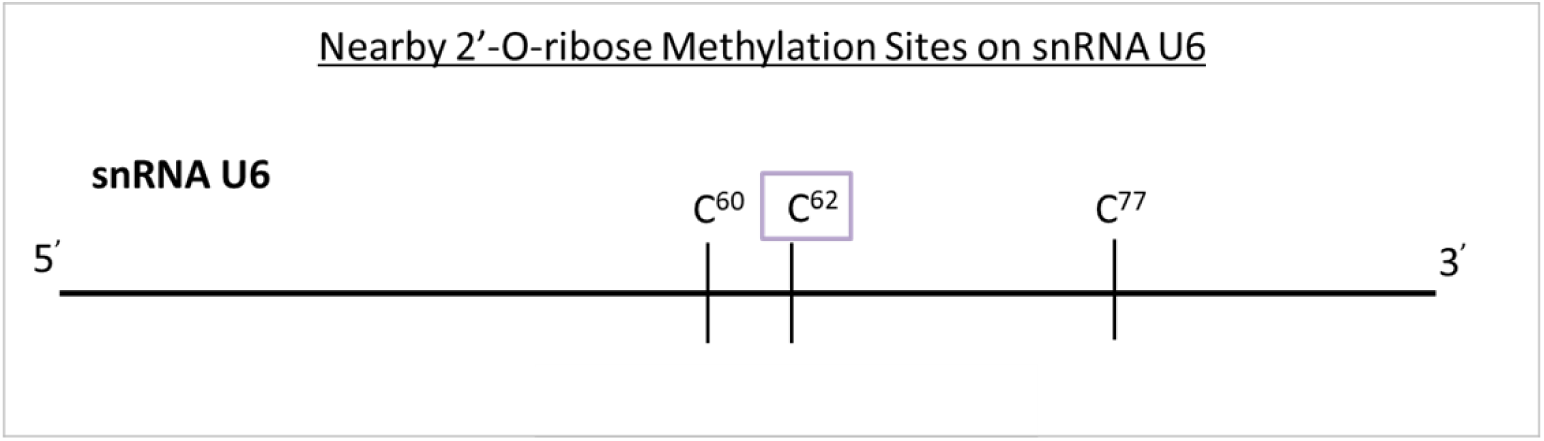
2’-O-ribose methylation sites on spliceosomal RNA U6 in proximity of C^62^. There are two nearby sites of methylation on U6, one two nucleotides upstream (C^60^) and one 15 nucleotides downstream (C^77^) of the target nucleotide of SNORD94 (C^62^) [18].

In the same snoRNA expression study, it was also shown that U6 had reduced expression by 3.2-fold in all 16 TOF samples as compared to control [3]. Analysis of splicing variation showed a substantial disturbance in splicing of genes known to be critical for heart formation, and many of these were in common with fetal patterns [3, 8]. This dramatic shift in ncRNA expression and splicing patterns may have resulted from failure of regulatory programs to progress properly during fetal heart development, perhaps contributing to a breakdown of spatially and temporally correct gene expression and splicing of mRNA contributing to the heart defect [3]. Here we quantified methylation at C^62^ on snRNA U6 to establish a link between levels of SNORD94 and levels of C^62^ methylation. These data may help decipher the underlying biochemistry essential for correct spatial and temporal gene expression patterns in key regulatory pathways in a developing vertebrate heart.

## Results

To quantify methylation at C^62^ in our samples, we followed the approach used by Dong et al. to quantify 2’-O-methylated cytosine [17]. This protocol was based on data from previous rRNA studies that showed that reverse transcription terminates at 2’-O-methylated cytosines in dNTP concentrations are limiting [20]. For each RNA sample, 50ng total RNA was reverse transcribed using two different dNTP concentrations (1mM or 1μM). The standard concentration (1mM) of dNTP allowed unimpeded reverse transcription as was previously shown [20]. However, the low concentration (1μM) causes stalling of reverse transcription at the methylated nucleotides resulting in shortened cDNA fragments [17, 20]. Quantitative Real-Time PCR with appropriate primers (see Methods) was used to quantify the long fragments between the RT reactions with normal and low dNTP concentrations. Thus, greater methylation (mC^62^) causes increased stalling of the reverse transcriptase resulting in less long fragments that leads to a relatively greater ΔC_t_ (Figure 2) when comparing samples of high versus low methylation.

**Figure 2.**
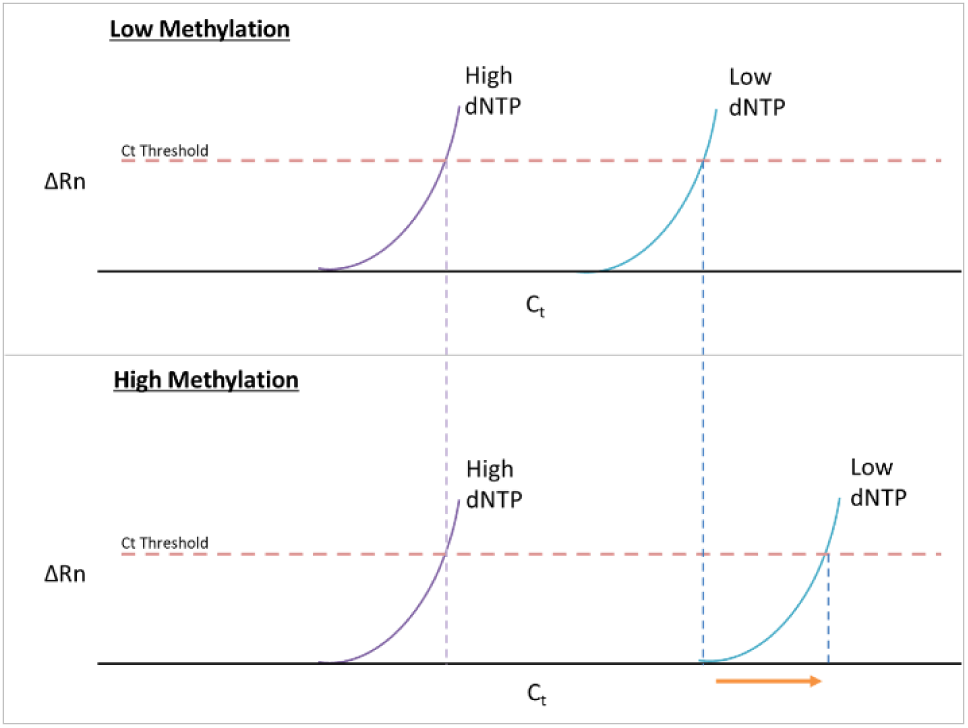
Idealized plots of qRT-PCR results for a sample with low mC^62^ vs high mC^62^. (TOP) An idealized plot qRT-PCR results for a sample with low mC^62^. There is a loss of long fragment in low dNTP concentration shown by the change in C_t_ between high and low dNTP concentrations. (BOTTOM) An idealized plot of qRT-PCR results for a sample with high mC^62^. The change in long fragment quantity is larger as compared to the low methylation group (Arrow). The long fragment in low dNTP concentration is reduced in quantity when methylation is high.

We validated the technique for quantifying methylated cytosine by using synthesized sequences corresponding to the sequence from human U6. RNA oligos were custom synthesized by IDT (Integrated DNA Technologies; Coralville, Iowa) and corresponded to the sequence of whole human U6. One oligonucleotide was methylated at C^62^ (mC^62^) and the other was unmethylated. We used 0.1ng RNA for each RT reaction (instead of 50ng as with the total RNA samples). We compared the cycle change for long fragments in low vs normal dNTP concentrations (ΔC_t_ long fragments) and found that the difference is significantly greater when methylation is higher. We also found that there exists a nonzero ΔC_t_ when no methylation is present, showing there is some loss in overall RT efficiency in low dNTP concentration.

Figure 3 shows the observed ΔC_t_ value for the difference in long fragment quantity between normal and low dNTP concentrations with synthesized fragment with either 100% methylation at C^62^ on U6 or 0% methylation.

**Figure 3.**
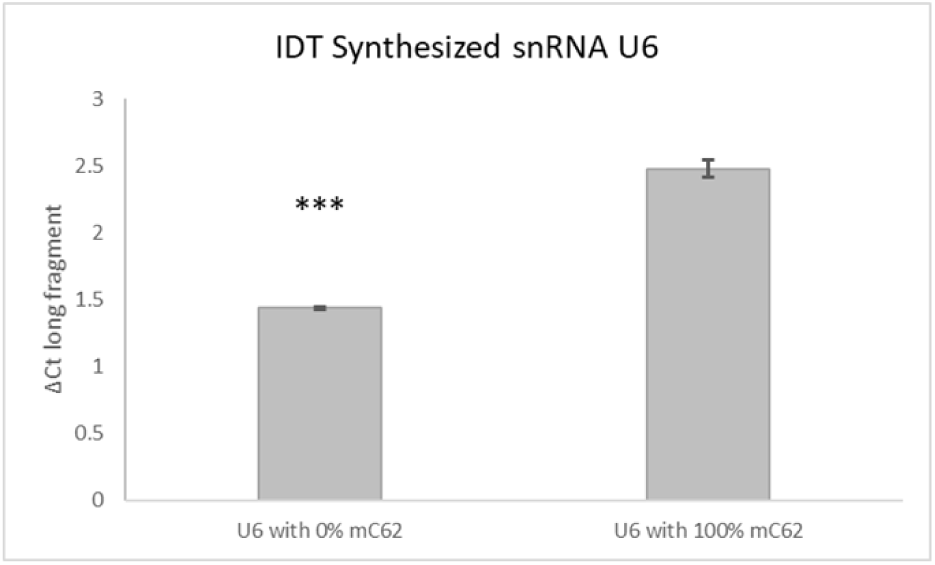
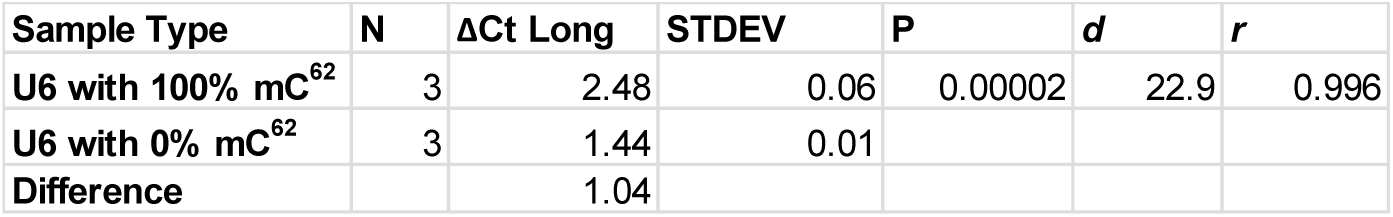
Validation of Technique with IDT Synthesized whole snRNA U6 with mC^62^ vs with no methylation. Treatment with low dNTP concentration results in a corresponding change in R based on level of methylation present in a sample (N=3, effect-size r = 0.98). (N=number of samples; STDEV=standard deviation; *d*=Cohen’s d; *r* =effect-size r; ***= p<.001).

We analyzed the data from all groups using t tests for the difference between groups. Because of the small sample sizes in each comparison group, we also calculated the Cohen’s *d* and effect-size *r* for each group, which are measures for the power of the difference in treatment groups with small sample size. [21, 22]. Larger Cohen’s *d* (*d*>0.8) is considered a “large effect” of treatment on the experimental group as compared to control, and *r* is another effect-size measure that represents the percentile of the treated group in relation to the control group [22]. Larger *d* value and *r* values correspond to greater effect-size of the treatment as compared to control [22].

After establishing the validity of the process, we proceeded to evaluate methylation of C^62^ in RNA isolated from tissues and cells. We compared the amount of mC^62^ in the right ventricular tissue of infants with TOF compared to right ventricular tissue from infants with normally developing hearts. We found that the patients with TOF had a significantly reduced amount of mC^62^ in right ventricle tissue as compared to age-matched normally developing infants (P<0.05) (Figure 4).

**Figure 4.**
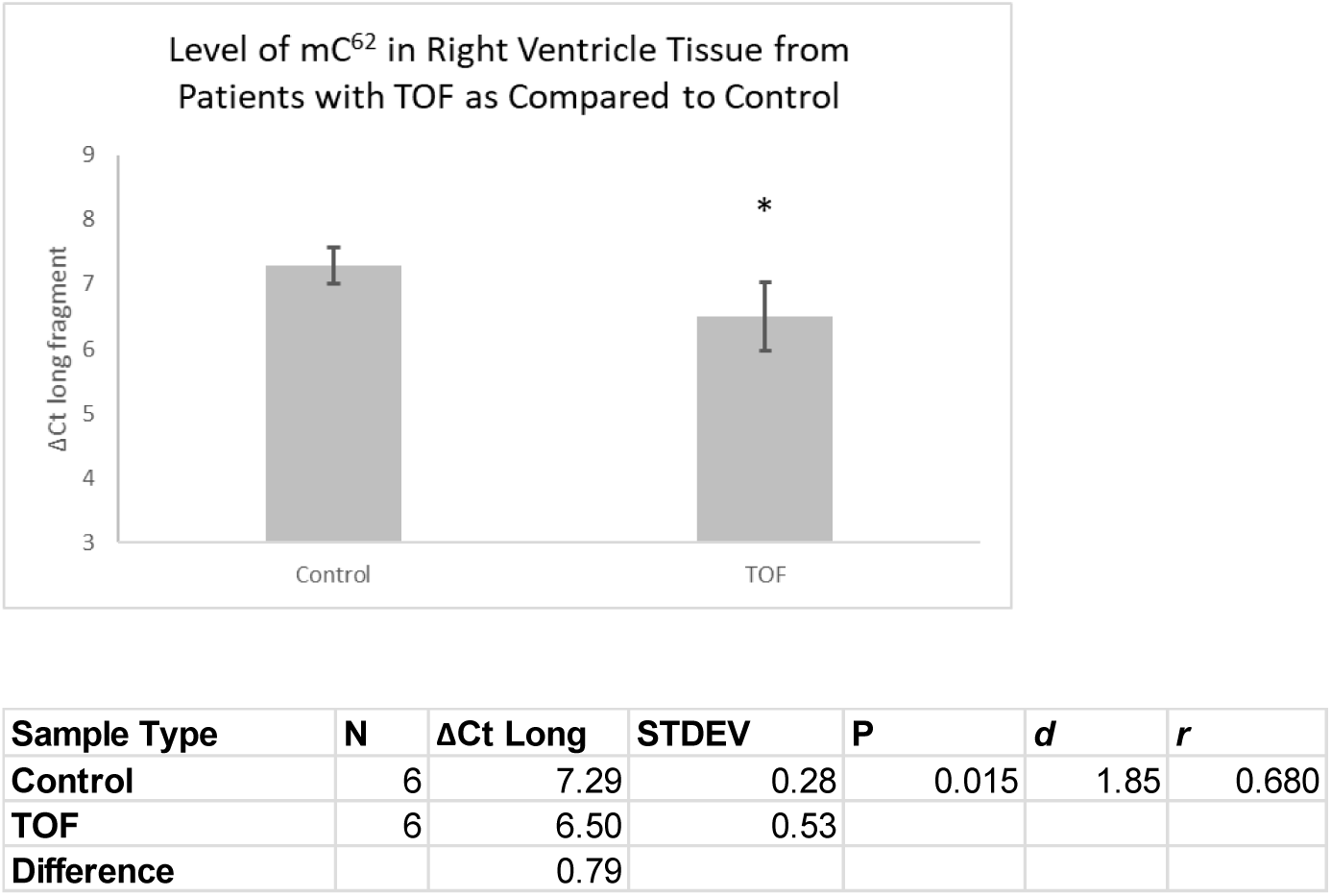
Methylation quantity in Right Ventricle Tissue from Patients with TOF as Compared to Age Matched Controls. Significant reduction in methylation in TOF tissue. (N=number of samples; STDEV=standard deviation; *d*=Cohen’s d; *r*=effect-size r; *= p<0.05)

In our *in vitro* assessment of altered levels of SNORD94, we found that knockdown of SNORD94 in primary cardiomyocytes derived from a normally developing heart caused a significant decrease in methylation at C^62^ (P<0.05) (Figure 5). Finally, we found that over expression of SNORD94 in primary cardiomyocytes from a patient with TOF showed an increased level of methylation at C^62^ on U6, however this difference was not significant at this sample size (P=0.12) (Figure 5)^1^.

**Figure 5.**
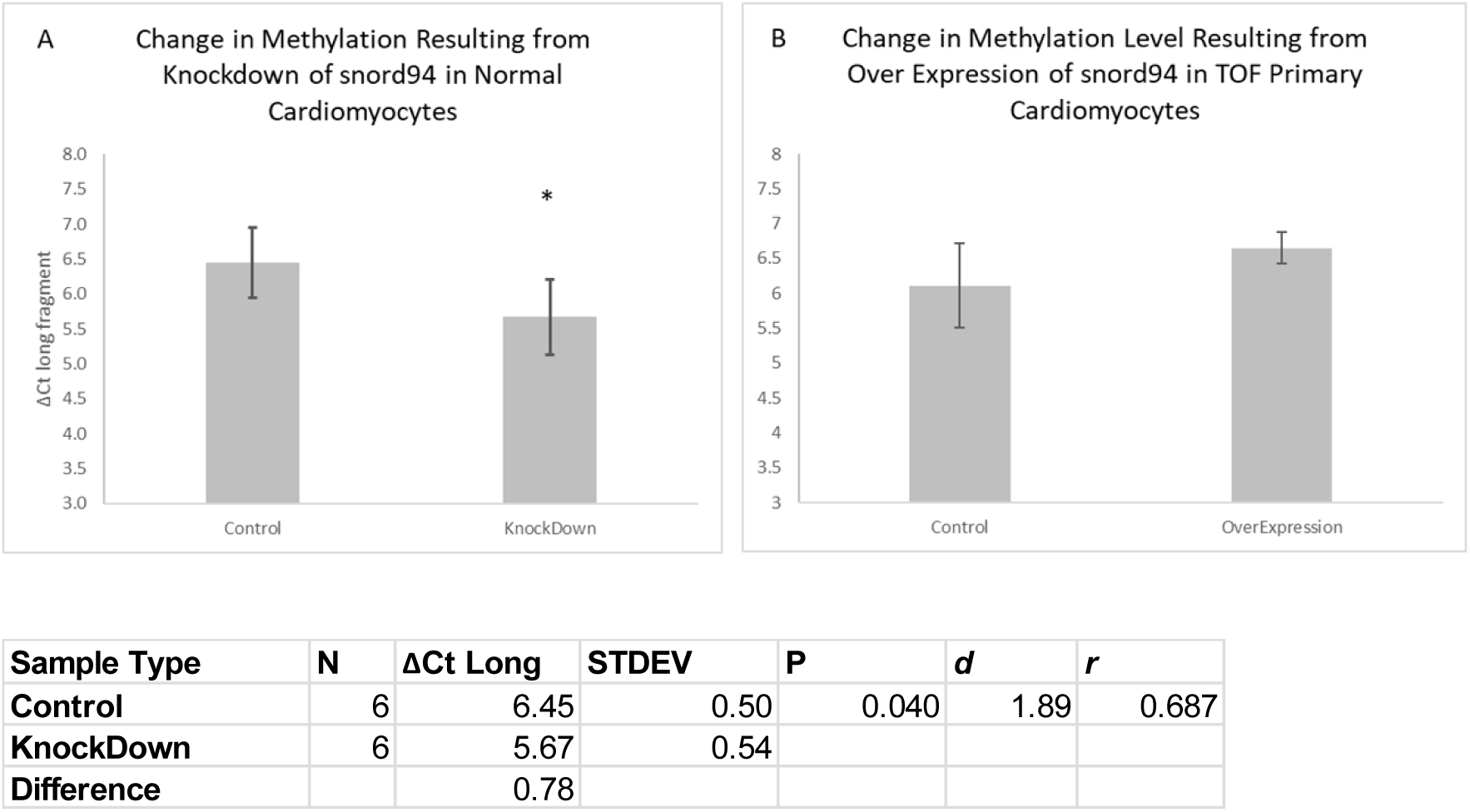
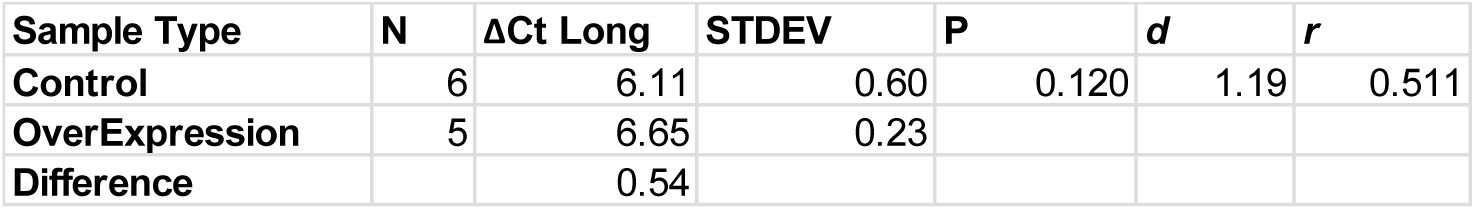
Changing expression of SNORD94 causes changes in methylation levels. A) Significant negative responses in methylation level according to knockdown of SNORD94. B) Non-significant positive responses in methylation level according to overexpression of SNORD94. (N=number of samples; STDEV=standard deviation; *d*=Cohen’s d; *r*=effect-size r; *= p<0.05).

Of note, there are other known sites of 2’-O-ribose methylation within spliceosomal RNA U6 (Figures 1 and 6). These methylation sites may have contributed somewhat to the loss of long fragment in low dNTP concentration for the total RNA samples (these methylation sites are not included in IDT synthesized U6) as these methylation sites may have also caused RT stops. However, the change in SNORD94 expression was the only difference in the knockdown and over expression in the cell culture experiments. This supports our contention that mC^62^ contributes the most to the difference in long fragment between groups with higher known mC^62^, as it did with the IDT synthesized oligopeptide validation. The loss of long fragment seen is likely slightly affected by methylation at C^77^ and C^60^ because of the proximity of C^60^ (although upstream) to our nucleotide of interest and the downstream location of C^77^. More research is needed to systematically determine the contribution of each methylation point to the overall loss of long fragment, with particular regard to C^60^ as the scaRNA that modifies it does have reduced expression in RV tissue from infants with TOF[8]. However, it appears that the influence on long fragment quantity of any methylation changes at C^60^ is small because we observed significant changes in long fragment loss when only SNORD94 expression was changed *in vitro*.

**Figure 6.**
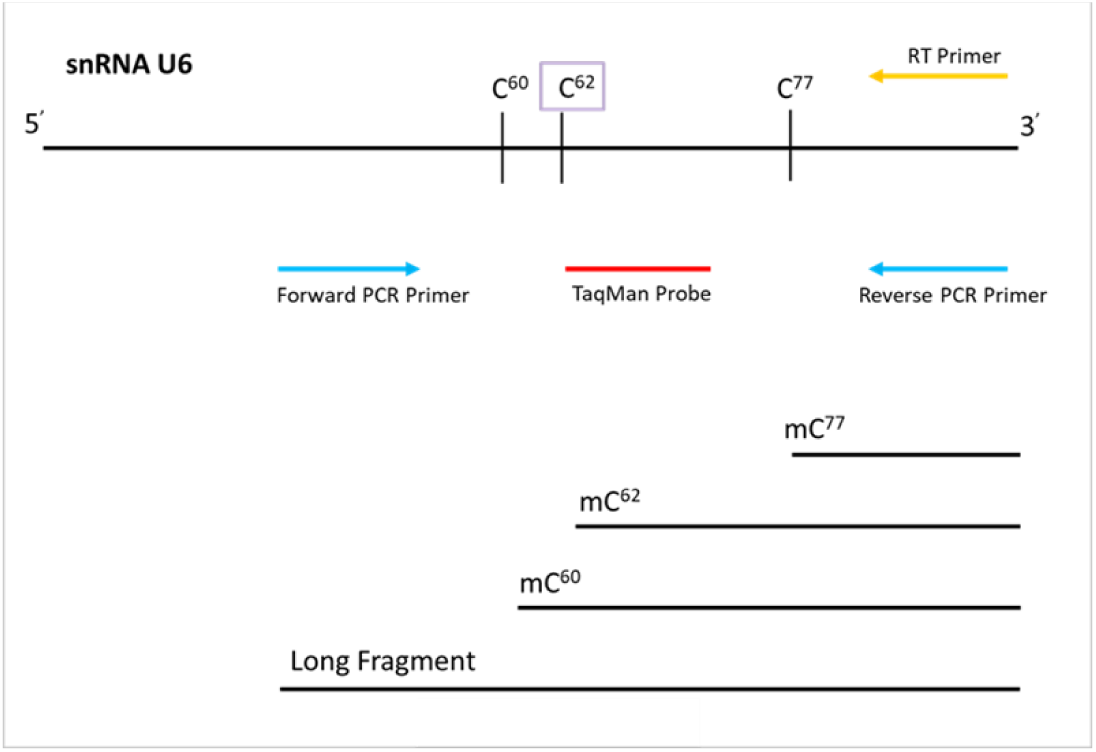
Schematic of Primer Design used in RT and qRT-PCR with potential resulting RT fragments in low dNTP concentration. The reverse primer used for RT (yellow) was a specific stem-loop primer binding to the 3’ poly-A tail end of U6, custom built for use with the PCR primer pairs and TaqMan probes from the same kit. The PCR primer pair (blue) used to detect either the long or the short fragments in qRT-PCR is also shown, with the TaqMan probe (red) positioned between the forward and reverse PCR primers. Primer locations are approximate based on our specifications for Thermo Fisher in designing the custom kit. The potential RT fragments produced in low dNTP concentrations based on locations of 2’-O-ribose methylations are also shown. During qRT-PCR amplification, only the long fragment is amplified.

## Discussion

Alternative splicing has a significant impact on developmental pathways in multiple organ systems, including the heart. We have shown that methylation at C^62^ on U6 is reduced in right ventricle tissue of children with TOF as compared to control. We have also confirmed that changing levels of SNORD94 in cell culture produces a corresponding change in methylation at C^62^ within spliceosomal RNA U6. These findings taken together suggest a dysfunction in an important regulatory pathway that is responsive to scaRNA expression levels. Directly stated, our accumulated results suggest that dysregulated scaRNA levels contribute to defective heart development by disturbing proper alternative splicing of mRNAs. This study focused on one specific scaRNA producing one biochemical modification, but more research is necessary to determine the interplay of multiple scaRNAs on spliceosomal integrity.

Our previous studies provided evidence that modest dysregulation of scaRNA expression impacted mRNA alternative splicing and development [3, 6, 8, 23]. Here we present evidence that there is a direct connection between the expression level of a single scaRNA and the level of methylation at its target nucleotide. This provides supporting evidence that the level of scaRNA expression directly influences alternative splicing through its biochemical function. Collectively these data suggest that there is likely a biological threshold of accumulated change in scaRNA expression that may be important contributors to heart defects.

## Materials and methods

### Subjects and tissue acquisition

Our subjects included 16 nondysmorphic infants (eight male, eight female) with idiopathic TOF but without chromosome abnormalities (22q11.2 deletion) [3, 23]. Comparison tissues from five (two male, three female) normally developing infants were obtained from LifeNet Health (https://www.lifenethealth.org, Virginia Beach, VA), a non-profit regenerative medicine company that provides bio-implants and organs for transplantation [24]. The control subjects were matched for age to the study population and all control subjects expired due to non-cardiac related causes. All donor tissue was de-identified, no donor confidential information was disclosed, and consent was obtained to use the tissue for research [23]. Tissue samples from the study subjects were collected at the time of surgical correction of TOF. Informed consent was obtained from a parent or legal guardian after reviewing the consent document and having their questions answered (Children’s Mercy Hospital, IRB # 11120627).

Primary cell cultures were derived from right ventricle (RV) tissue of infants with TOF. The RV tissue was immediately immersed in DMEM (Invitrogen/Gibco, Grand Island, NY) plus 10% fetal calf serum (Sigma/Safc, St. Louis, MO) and 1% penn/strep (Gibco). The tissue was minced and most of the media was removed, leaving only enough to keep the tissue from drying out. After 24 hours, additional media was added, and cells were growing robustly after 3 to 4 days. Media was exchanged every 48 hours. In addition, we obtained a primary neonatal cardiomyocyte cell culture derived from normally developing human neonatal cardiac tissue from Celprogen (San Pedro, CA Cat#36044-21). These cells were also grown in DMEM plus 10% fetal calf serum and 1% penn/strep.

### Transfection of scaRNA plasmids into primary cells

The SNORD94 expression vector used in this study was that which was used in Patil et al, 2015 and was a generous gift from Dr. Tamas Kiss [8]. The scaRNA was cloned into an intron sequence between hemoglobin exons 3 and 4 so that they would be correctly processed in vivo, and expression was driven with the CMV promoter. The scaRNAs were transfected into the primary cell lines derived from infants with TOF according to the manufacturer’s protocol and as done previously [8]. Briefly, 2 µg of plasmid DNA was diluted in 200 µL of serum free media and added to 2 µL of the Poly Magnetofectant (a magnetic nanoparticle transfection reagent; Oz Bioscience, France), vortexed, and incubated for 20 min at room temperature. The transfection mixture was added dropwise to cells in 1.8 ml of media containing 10% serum in a single well of a 6 well plate. The culture plate was set on top of a plate magnet (Oz Biosciences) for 20 min and returned to the incubator. After 72 hours, the cells were trypsinized, pelleted and stored at −80 °C until processed for RNA extraction.

### scaRNA knockdown

We used an antisense LNA oligo (locked nucleic acid oligos, Exiqon Life Sciences, Woburn MA) to suppress the SNORD94 in primary cardiomyocytes as has been done previously in immortalized cell cultures [8, 25, 26]. Briefly, the LNA oligo protocol is as follows, 50 µM LNA oligo in 100 µL serum free media is mixed with 12 µL HiPerfect transfection reagent (Qiagen, Valencia, CA) and incubated for 20 min at room temperature. The transfection mixture was added to cells in 2.3 mL of media with 10% serum in a single well of a 6 well cell culture plate. After 48 hours, the cells were pelleted and stored at −80 °C until processing. The LNA oligo used in this study was previously found effective by empirical process [8].

### RNA isolation

RNA from tissue or cell culture was extracted from the whole cell pellet using a mirVana total RNA isolation kit (Invitrogen) according to the manufacturer’s instruction.

### Quantification of 2’-O-ribose methylation

It has been previously shown that reducing concentration of dNTP to 4μM or below in the cDNA synthesis reaction can induce a stop at a 2’-O-ribose methylated nucleotide on RNA [17, 20]. We used the same approach but modified the protocol for use with custom TaqMan probes for high specificity in quantitative Real-Time PCR. Reverse transcription and quantitative PCR were done for each RNA sample using a custom TaqMan Small RNA Assay Kit (Applied Biosystems, catalog #4398987) for the long fragment of U6 created when no methylation is present (ID #CTMFW2X), and no stop occurs. This kit contains a stem-loop reverse transcription primer and a mix of forward and reverse PCR primers with TaqMan probes for use with PCR (shown in Figure 6). The custom TaqMan Small RNA Assay Kit was used according to manufacturer’s protocol except for two notable exceptions: reducing dNTP concentration to 1µM in the “low dNTP concentration” group as compared to 1mM concentration in the “normal dNTP concentration” group and using 50ng total RNA per reaction rather than the recommended 10ng because the reduced dNTP concentration also tends to cause a reduction in overall reverse transcription. In total, for each RNA sample there were 2 different reactions created using either the RT kit, and either normal or low concentration of dNTP. RNA was split from the same aliquot for each RT reaction. Quantitative Real-Time PCR was then performed using the ViiA 7 from Applied Biosystems using standard conditions and an annealing temperature of 60°C.

### Statistical Calculations

The RT reaction stalls at 2’-O-ribose methylated cytosines (mC) when dNTPs are in low concentration. Thus, the loss of the long fragment in low as compared to normal dNTP concentrations will reflect the amount of mC62. However, there is some reduction in efficiency of reverse transcription in the presence of low dNTP concentration, so there will be a nonzero change in long fragment quantity in the absence of methylation. We used the difference in cycle number (ΔCt) as a measure of methylation; a greater ΔCt represents a larger amount of mC62 due to a greater loss of long fragment in low dNTP concentration as compared to high concentration. An increased level of methylation in an RNA sample results in a reduced amount of long fragments produced in low dNTP concentration during the RT reactions. These changes during RT will result in a larger ΔCt in qRT-PCR.

## Acknowledgements

A special thanks to Marissa Roffler, PhD (KCU), for her statistical advice.

## Supporting information

S1 files

Data tables

## Footnotes

1 Within the Over Expression experiment, there were one over expression sample that were excluded from the data analysis due to RNA degradation shown on qRT-PCR when tested for SNORD94 expression.

